# Montage electron tomography of vitrified specimens

**DOI:** 10.1101/2021.11.02.466666

**Authors:** Ariana Peck, Stephen D. Carter, Huanghao Mai, Songye Chen, Alister Burt, Grant J. Jensen

## Abstract

Cryo-electron tomography provides detailed views of macromolecules *in situ*. However, imaging a large field of view to provide more cellular context requires reducing magnification during data collection, which in turn restricts the resolution. To circumvent this trade-off between field of view and resolution, we have developed a montage data collection scheme that uniformly distributes the dose throughout the specimen. In this approach, sets of slightly overlapping circular tiles are collected at high magnification and stitched to form a composite projection image at each tilt angle. These montage tilt-series are then reconstructed into massive tomograms with a small pixel size but a large field of view. For proof-of-principle, we applied this method to the thin edge of HeLa cells. Thon rings to better than 15 Å were observed in the montaged tilt-series, and diverse cellular features were evident in the resulting tomograms. These results indicate that the additional dose required by this technique is not prohibitive to performing structural analysis to intermediate resolution across a large field of view. We anticipate that montage tomography will prove particularly useful for lamellae, increase the likelihood of imaging rare cellular events, and facilitate visual proteomics.

## I. Introduction

Cryo-electron tomography (cryo-ET) is a powerful tool for studying macromolecular structures in the near-native context of frozen-hydrated cells, unperturbed by stains or fixatives [1]. In this cryo-electron microscopy (cryo-EM) technique, a series of projection images is recorded as a vitrified specimen is tilted in an electron microscope. The resulting tilt-series is reconstructed into a tomogram, or volumetric map of the specimen’s electrostatic potential. Cryo-ET has revealed important details of cellular ultrastructure, and subtomogram averaging algorithms enable determining the structures of macromolecular complexes at a resolution of 0.4-4 nm [1]. Recent technical advances have extended the high-resolution potential of this technique, yielding subtomogram averages of purified HIV-1 Gag particles to 3.1 Å and *in situ* ribosomes to 3.7 Å [2, 3]. However, data must be collected at high magnification to retain this high-resolution signal. The trade-off is a smaller field of view, limiting the region that can be imaged to a tiny fraction of a cell.

In principle, montage tomography could permit imaging a large field of view without sacrificing high-resolution details. For montage data collection, the beam is tiled across a specimen and the recorded images are computationally stitched together during reconstruction [4]. To date, montage tomography has only been performed on resin-embedded samples, which resist radiation damage but suffer from artifacts induced by chemical fixation [5–7]. By contrast, cryo-preserved specimens can tolerate only a limited electron dose before being destroyed [8]. Historically this dose sensitivity has been considered prohibitive to collecting montage data from vitrified samples, as portions of the sample must be exposed multiple times to facilitate stitching images during reconstruction.

Recent technical developments, however, motivate revisiting the potential of montage cryo-tomography. First, the highly stable optics of modern microscopes enable precise control over the region being exposed [9]. Such precision is critical both to prevent gaps between neighboring images and to avoid accidentally enlarging the overlap region. Second, the ability to collect data using a circular beam with fringe-free illumination allows for more efficient tiling strategies and reduces loss of information due to corruption by Fresnel fringes [10, 11]. Third, the increased sensitivity of modern detectors permits decreasing the exposure while achieving the same signal-to-noise ratio [12–15]. Fourth, focused ion beam (FIB) milling has significantly expanded the range of specimens that can be studied by cryo-ET but is highly time-consuming, motivating efforts to image as much of the lamellae as possible [16, 17]. In addition, there is growing recognition in the field that radiation damage is a progressive phenomenon. As a result, important features of cellular biology remain observable even after receiving what was previously considered an intolerably high dose for vitrified samples.

Montage tomography of cryo-preserved specimens would further the potential of cryo-ET by increasing the likelihood of imaging transient events and providing significantly more cellular context for macromolecules of interest. Here we use simulations to optimize a montage data collection scheme and develop strategies to stitch tiles at each tilt angle into a composite projection image. We then harness modern cryo-ET algorithms to reconstruct tomograms from these montage tilt-series with a small pixel size but a large field of view. To demonstrate proof-of-principle, we applied this technique to the thin edges of HeLa cells. Thon rings to better than 15 Å were observed at both low and intermediate tilt angles of the montage tilt-series. The reconstructed tomograms spanned a 3.3 *μ*m^2^ field of view and contained diverse cellular features, including mitochondria, multilammelar vesicles, and microtubules. These results indicate that despite the additional dose required by this method, montage tomography enables capturing large fields of view for structural analysis at the intermediate resolutions typical of cryo-ET data.

## II. Optimization of tiling strategies

Given the sensitivity of biological samples to radiation damage, the success of montage tomography depends on efficiently distributing the total exposure both at each tilt angle and across the full tilt-series. The former is readily addressed for a circular beam: the optimal strategy to pack circles in a plane uses a hexagonal tiling scheme, in which circles are centered on the vertices of a regular hexagonal grid and three neighboring circles intersect at a point (Fig. 1A) [18]. The question then remains how to displace these hexagonally-packed circular tiles between tilt angles to most uniformly spread the dose across the tilt-series. Applying global translations and rotations to the hexagonal array of circles between tilt angles changes which regions of the sample lie in an overlap region at each tilt angle, thereby reducing the amount of sample that receives excess dose (Fig. 1B-C).

**FIG. 1.**
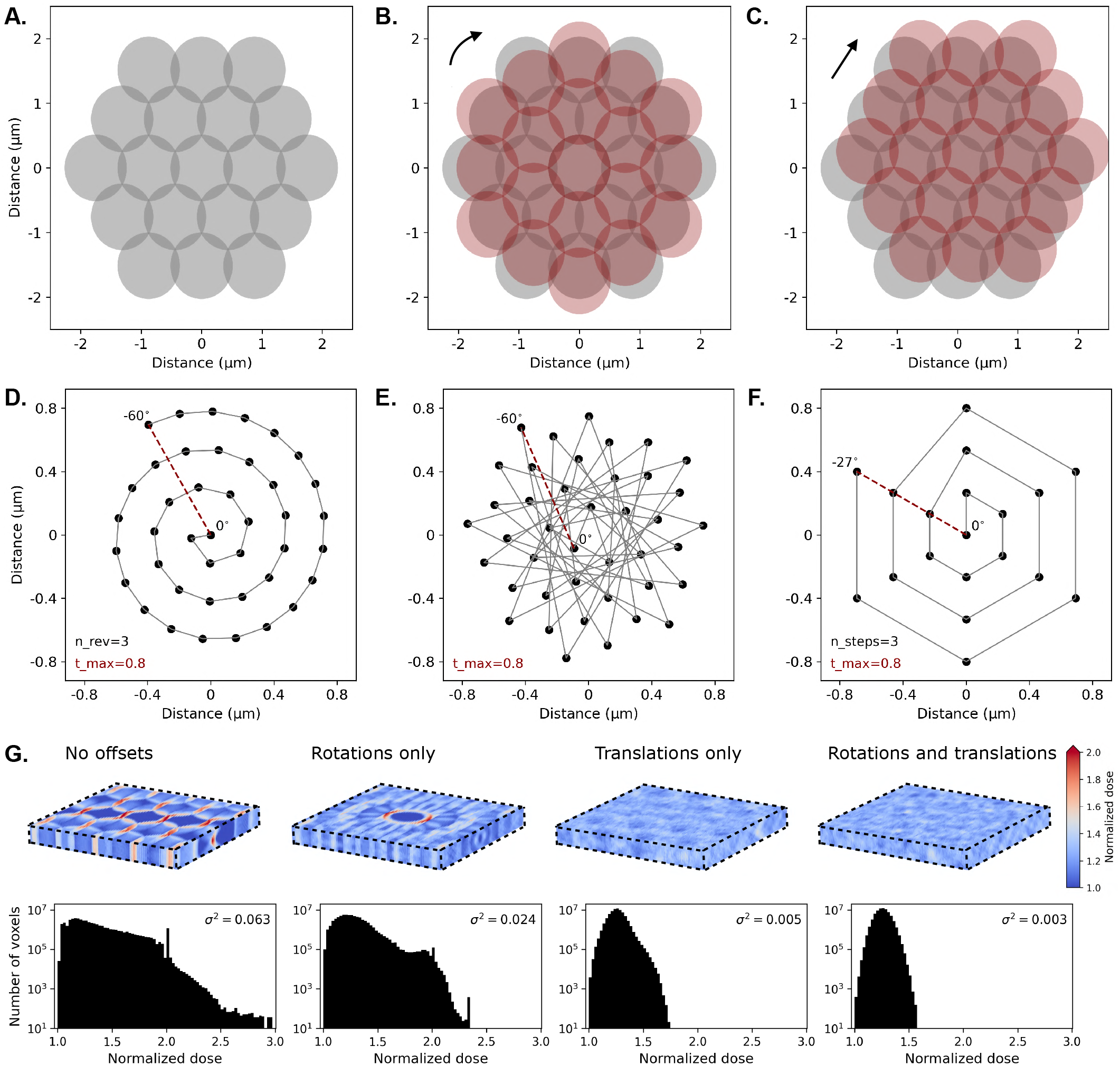
Optimization of a montage tiling strategy. (A) At each tilt angle, a hexagonally-packed set of circular tiles is imaged. Applying a global (B) rotation or (C) translation to the tiles between tilt angles changes which regions of the specimen lie in an overlap region to more uniformly spread the dose. To systematically introduce translational offsets between tilt angles, the global displacements applied to the tiles between tilt angles followed one of three spiral patterns: (D) an Archimedean spiral, (E) a sunflower spiral, or (F) a “snowflake” spiral. The positions of the central tile are indicated by black dots and spiral outwards during the course of the tilt-series. For the snowflake spiral, the pattern is repeated starting from the center if the outermost position is reached before the final tilt angle. One of the tunable parameters for all three spiral patterns is the maximum translation of the central tile (Emax); a second parameter for the Archimedean and snowflake patterns is the number of revolutions (n_rev or n_steps). (G) The dose distributions received by voxels of a discretized specimen during a simulated tilt-series are compared for four tiling strategies. The spatial distribution of the dose is mapped on the specimen (upper). Each tiling strategy was scored by the variance (*σ*^2^) of the dose distribution, which is noted at the upper right of each histogram (lower). For the “no offsets” strategy, the hexagonally-packed tiles were rotated by 10° relative to the plane of the detector but no offsets were applied to tile positions between tilt angles. For the “rotations only” strategy, a 20° clockwise rotation was applied to all tiles between tilt angles. For the “translations only” strategy, tile positions were translated along an Archimedean spiral with three revolutions and a maximum translation of 80% of the beam radius for each tile. The best-ranked strategy (right) applies both of these translational and rotational offsets to efficiently distribute the dose.

Since *a priori* it is unclear which combination of offsets would most efficiently distribute dose, we used simulations to characterize hundreds of different tiling strategies. These simulations used a right-handed coordinate system with the detector oriented in the *xy* plane and the incoming electron beam directed along the *z*-axis (Fig. S1). A rectangular specimen with a width of 3,420 nm and depth of 400 nm was discretized into 4 nm cubic voxels and tilted about the *x*-axis following a standard dose-symmetric tilt-scheme [19]. At each tilt angle, a 1 *μ*m diameter beam was used to illuminate a set of hexagonally-packed circular tiles. Even with fringe-free illumination, residual fringes were observed to affect up to 2% of the outer edge of each tile (see below). To prevent gaps in the montage after discarding this corrupted region, the overlap between tiles was increased relative to optimal hexagonal packing such that a small fraction of voxels was exposed up to three times at any tilt angle. We then computed the accumulated dose received by each voxel of the specimen under tiling strategies that differed in the translational and rotational offsets applied between tilt angles.

To systematically introduce translational offsets, we examined three basic spiral patterns: an Archimedean spiral, in which adjacent points are equidistant along the curve of the spiral (Fig. 1D); a sunflower or Fibonacci spiral, in which points are distributed in concentric shells of equal area, and successive points are placed in the largest angular gap between previous points [20] (Fig. 1E); and a “snowflake” spiral, in which points are positioned on the vertices of concentric hexagons that mirror the 6-fold symmetry of the packed circular tiles (Fig. 1F). The positions of all tiles were uniformly shifted between successive tilt angles to follow the path of the spiral while maintaining hexagonal packing at each tilt angle. For all three spiral types, one of the adjustable parameters was the maximum translation permitted for each tile across the entire tilt-series (Fig. S2A); this was capped at a distance of one beam radius to ensure that all but the outermost tiles remained fully in the field of view. The second translational parameter was the number of revolutions or radial steps respectively for the Archimedean spiral and snowflake pattern (Fig. S2B). For the sunflower pattern, positions were dictated exclusively by the number of points and maximum translation. A third translational parameter was an optional scaling of the *x*-axis displacements by 1/cos *α*, where *α* is the tilt angle (Fig. S2C). This scaling mimicked the *y*-axis elongation of circular tiles into ellipses at high tilt angles (Fig. S1), with the intent of preserving efficient circular symmetry.

In addition to translational offsets, rotational offsets were systematically introduced by varying three parameters. The first parameter was the starting angle, which dictated the initial orientation of the hexagonal array of circular tiles in the plane of the detector. The second parameter was the rotational step size, which determined the magnitude of the global rotation applied to the hexagonally-packed tiles between each tilt angle. Third, this global rotation was either applied continuously or in an alternating fashion. For the continuous scheme, the hexagonally-arranged tiles were rotated by the same amount and in the same direction between each tilt angle. For the alternating scheme, global counterclockwise and clockwise rotations of the same rotation step size were applied between successive tilt angles.

In total we simulated 576 snowflake, 546 spiral, and 286 sunflower patterns by systematically varying the parameters described above. Each pattern was scored by the variance of the distributed dose, with superior patterns characterized by low variance (Fig. 1G). Across all spiral types, we found that translational offsets were more critical than rotational offsets to uniformly spread the dose throughout the specimen (Figs. 1G, S3). Although the top-ranked variant for the (Archimedean) spiral, sunflower, and snowflake patterns achieved similar scores, we found that variants of the first consistently scored well for translational offsets of the same magnitude (Fig. S3). All variants were also observed to perform similarly across different data collection schemes (Fig. S4). We thus chose the best-performing spiral variant for experimental data collection. This pattern was characterized by a maximum tile translation of 80% of the beam radius, 3 revolutions,no scaling of the *x*-axis translational component, a starting offset angle of 10° of the hexagonally-arranged tiles in the plane of the detector, and continuous rotations of 20° between tilt angles (Fig. 1G, right). These offsets reduced the variance of the distributed dose by 20-fold compared to the corresponding pattern without any offsets (*σ*^2^=0.003 versus *σ*^2^=0.063).

## III. Montage data collection and processing

### A. Sample preparation

HeLa cells (cell line 25) were cultured in DMEM without phenol red (Gibco) supplemented with 10% FBS, 100 U/mL penicillin, and 100 *μ*g/mL streptomycin at 37° C with 5% CO_2_. Cells were plated at a density of 2 × 10^5^ cells/mL on laminin-coated 200-mesh gold R3.5/1 London finder Quantifoil grids (Quantifoil Micro Tools) and incubated for ~12 hours. For a subset of grids, 3 *μ*L of a colloidal solution of 20 nm gold fiducials (Sigma-Aldrich) pre-treated with bovine serum albumin were applied immediately before vitrification. The grids were then plunge-frozen in a mixture of liquid ethane/propane using an FEI Vitrobot Mark IV [21] and stored in liquid nitrogen at ≤-150°C.

### B. Montage tilt-series collection

Montage tilt-series were collected on a Titan Krios G3i (Thermo Fisher Scientific) equipped with fringe-free illumination [10], a Gatan imaging filter, and a K3 Summit direct electron detector (Gatan). Data acquisition was performed using SerialEM in electron-counting mode [22]. Each tile was acquired using a circular beam of diameter 1.08 *μ*m, such that the beam spanned the short edge of the detector at a pixel size of 2.65 Å. A beam-centering step was performed after collecting each tile to reduce drift without applying additional exposure. At each tilt angle, 37 tiles were acquired: one central tile, with three surrounding rings of hexagonally-packed tiles. The beam coordinates were updated using image shifts to follow a spiral pattern over the course of the tilt-series as described in Section II. Data were collected using a grouped dose-symmetric tilt-scheme with a group size of 6°, 2° increments between tilt images, and a tilt-range of ±60° [23]. Applying the above data collection scheme at a constant defocus was predicted to yield a ~8 *μ*m defocus gradient at the highest tilt angles (Fig. 2A, left). To avoid this spread, we adjusted the defocus with which each tile was collected based on that tile’s estimated *z*-height in the microscope. Defocus values ranged from −5 to −11 *μ*m for different tilt-series, and a total dose of 60-106 e^−^/Å^2^ was used.

**FIG. 2.**
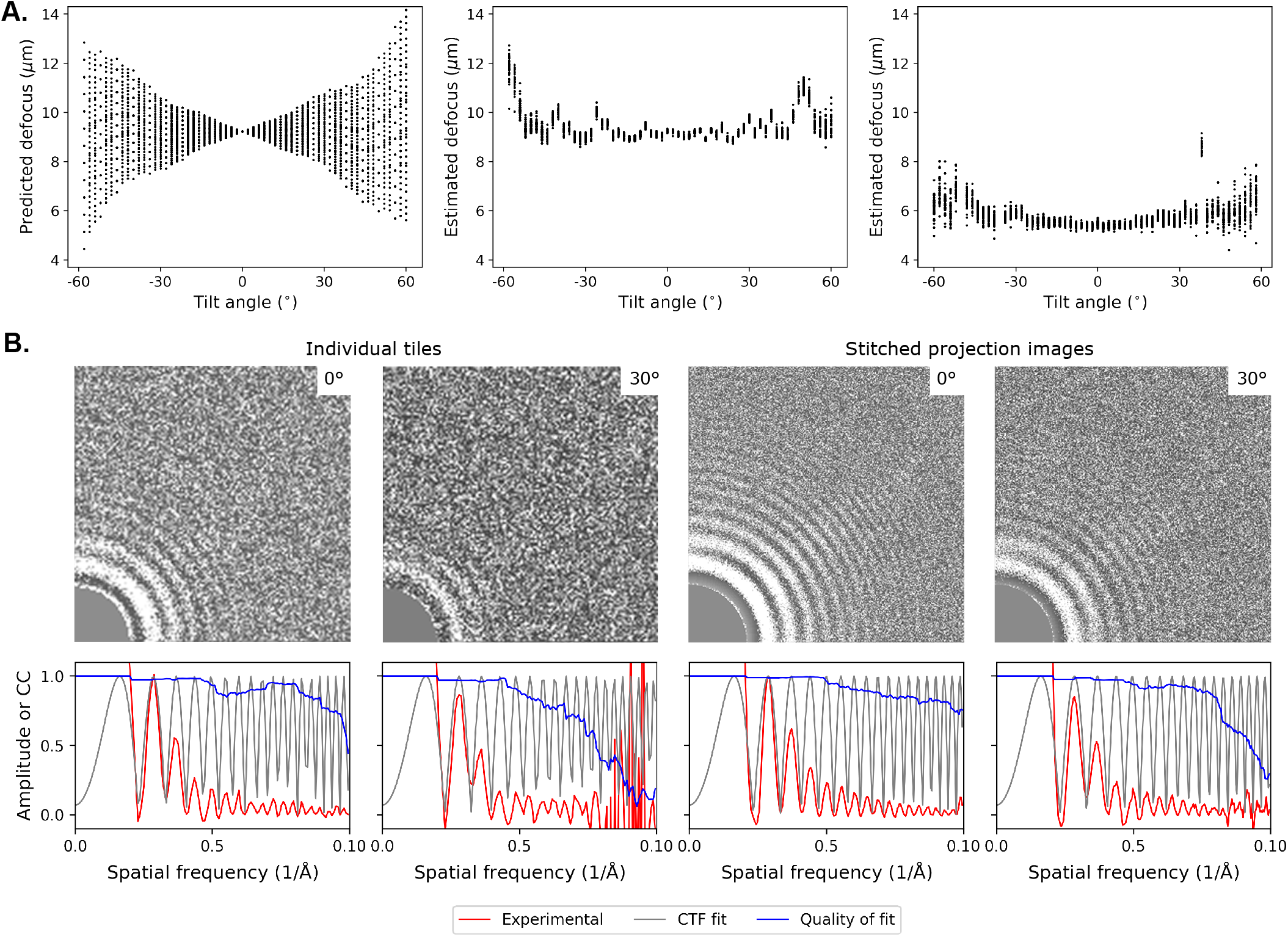
CTF estimation reveals a stable defocus throughout the tilt-series and Thon rings to better than 15 Å. (A) Defocus values were predicted based on the tiles’ estimated heights in the microscope and are shown as a function of tilt angle (left). By contrast, the per-tile defocus values estimated by CTFFIND4 showed a relatively stable defocus gradient, as plotted for two representative tilt-series (middle and right). This was accomplished by performing an autofocusing step prior to collecting each tile to compensate for the predicted defocus gradient across each tilt angle. (B) The 2D experimental spectrum (upper) and rotationally-averaged 1D CTF fits (lower) are shown for representative tiles (left) or CTF-uncorrected stitched projection images (right) at the indicated tilt angle.

### C. Tile pre-processing

Before stitching tiles into a composite image, Fresnel fringes and non-uniform illumination must be accounted for. Fringe-free illumination (FFI) reduces but does not entirely eliminate Fresnel fringes during image formation [10, 11]. Residual fringes are particularly evident when data are collected at a defocus and magnification typical for tomography, since current FFI set-ups are optimized for single-particle cryo-EM applications instead. The number of observed fringes depends on both defocus and the signal intensity of the specimen, so removing a fixed fraction of the outer edge of each tile does not adequately eliminate the fringe-contaminated region.

Given this variability, we developed the following heuristic approach to mask residual fringes. A 2-dimensional Gaussian bandpass filter was applied to each tile, using kernel sizes of 2 and 6 nm. In the filtered tile, a high-intensity ring that spanned the Fresnel fringes was observed at the edge of the illuminated region (Fig. S5A). A circle was fit to this ring of pixels using least-squares optimization, and the start of the Fresnel fringes was estimated as the ring’s inner radius. All pixels outside this radius were masked (Fig. S5B-E).

In addition to residual fringes, we observed a consistent reduction in radial intensity by ~15% between the center and edge of each tile (Fig. S5B). Left uncorrected, this would artificially depress the intensity of the overlap regions during stitching. We therefore applied a radial gain correction as follows. At each tilt angle, the tiles’ radial intensity profiles were normalized to a value of 1 in the central region of the tile and merged to generate a single intensity profile for the tilt angle. The resulting radial intensity profile was median-filtered and applied as a gain reference, with linear interpolation used to compute the correction factor at each pixel (Fig. S5B-E).

### D. Correcting for the contrast transfer function

Despite adjusting the focus on a per-tile basis to avoid an excessive defocus gradient across the tilt-series, some variation in defocus between tiles was expected due to microscope error. We therefore performed a contrast transfer function (CTF) correction on individual tiles. We used CTFFIND4 (version 4.1.13) to estimate each tile’s defocus [24] and *ctfphaseflip* to correct for the CTF [25]. The estimated defocus values confirmed that the per-tile focus adjustment compensated for the large changes in *z*-height due to montage collection and yielded a relatively stable defocus throughout the tilt-series (Fig. 2A). The high-resolution limit of detected Thon rings for most tiles ranged from 10-20 A, with better resolution at lower tilt angles as expected (Fig. 2B, left). To verify that the stitching procedure described below did not degrade resolution, CTFFIND4 was also applied to the CTF-uncorrected stitched images to assess the high-resolution limit of detectable Thon rings (Fig. 2B, right).

### E. Tile registration

We developed an image registration workflow tailored for montage cryo-tomograms collected using a circular beam. Current software for processing montage data is designed for resin-embedded specimens and unsuitable for two reasons. First, resin-embedded samples are typically acquired using the full area of the detector, yielding a large and rectangular overlap region between tiles [4]. By contrast, we employ a more efficient tiling strategy of hexagonally-tiled circular beams to overcome radiation sensitivity. This scheme results in lemon-shaped overlap regions that change positions and orientations relative to the specimen at each tilt angle (Fig. 1B-C). Second, the algorithms used to perform automated landmark extraction and alignment rely on high-contrast features that typify resin-embedded specimens but are absent in cryotomograms even when collected at high defocus [26].

To overcome low contrast, data collected with a pixel size of 2.65 Å were first binned to 10.6 Å. After applying a correction for uneven radial illumination, tiles were bandpass-filtered and masked to remove the unilluminated and Fresnel fringe-corrupted regions. For bandpass-filtering, we found that kernel sizes of roughly 6 and 19 nm enhanced features such as gold beads, membranes, and grid hole boundaries that serve as useful landmarks for image registration. These features were further selected for by thresholding; specifically, only pixels with intensities in the bottom 15th percentile for each tile (belonging to high contrast features) were retained.

Despite the stable optics of modern microscopes, some drift is expected during data collection. To refine the tile positions from the beam coordinates supplied to the microscope, we computed the translational shifts that maximized the normalized cross-correlation between pairs of overlapping tiles, *T* and *T*′, at each tilt angle:

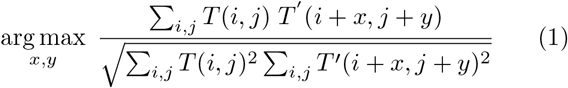

where the sum is over all pixels at coordinates (*i*, *j*) that are in register when *T*′ is translated by (*x*, *y*) while the position of *T* remains fixed. The initial search was performed in a box of length 30 nm centered on the beam coordinates used during data collection. If the translational shift that maximized the cross-correlation score was located at the edge of this box, the search box was re-centered to this position and another search was performed. This calculation yielded the relative positions for each pair of overlapping tiles.

The full mosaic for each tilt angle was then generated by fixing the central tile at the origin and determining the coordinates of the surrounding six tiles relative to this anchor tile (Fig. 1A). The position of each tile in the surrounding layer was estimated as the mean of all pairwise positions weighted by the cross-correlation scores between the tile of interest and each of its neighbors. Using the cross-correlation coefficients as weights ensured that overlap regions with the highest contrast features contributed most to tile positioning. The coordinates of the tiles in this first ring were then fixed. Tiles in each successive concentric ring were positioned using the same strategy, based on the consensus coordinates from pairwise registrations between neighboring tiles in the ring under consideration and anchor tiles in the previously fixed ring. Once the positions of the tiles in the outermost ring were determined, the full registration procedure was repeated using the optimized positions as the tiles’ starting coordinates. Tile registration was then performed on the unbinned data, using the optimized coordinates as the starting tile positions and a smaller search space.

Tiles were then stitched based on these optimized positions to generate a mosaic (Fig. 3, inset). For pixels lying in the overlap regions, the intensity values were selected from the tile whose center was nearest to the pixel. Continuous cellular features were observed in the overlap regions of the montage projection images, indicating minimal radiation damage and robust stitching (Figs. 3 and S6). Occasionally the combination of beam shift error and masking resulted in small gaps between tiles; these missing pixels were filled by randomly sampling intensity values in the surrounding region to prevent holes in the stitch. Each montage projection image was then shifted to compensate for the spiraling translational offsets applied between tilt angles, cropped to the maximal region imaged at all angles, and stacked to generate a tilt-series.

**FIG. 3.**
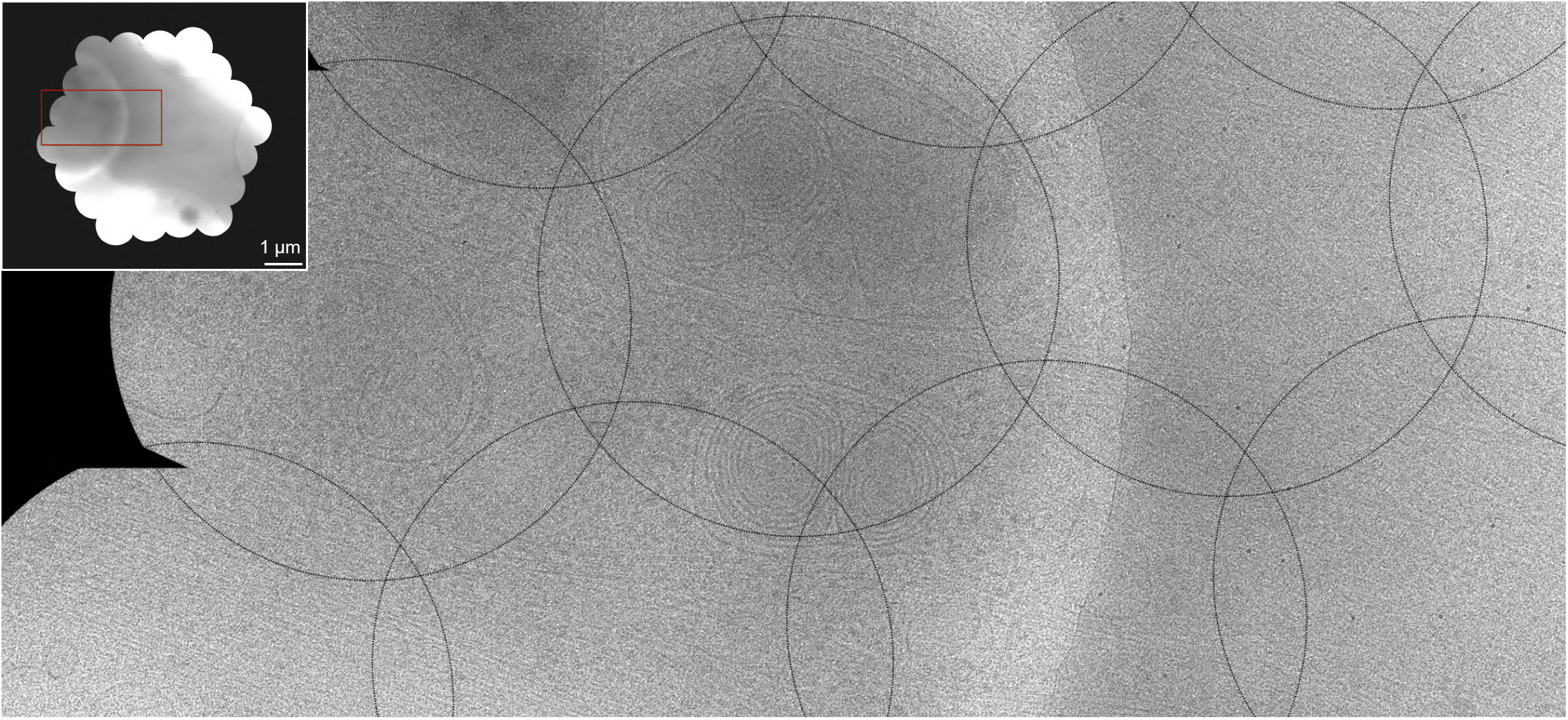
Continuity of cellular features in the overlap regions indicates successful stitching. The montaged projection image at 0° from a representative tilt-series is shown in the upper left inset. The region boxed in red is visualized at higher detail in the main image, with the boundaries of the circular tiles drawn in black. The clear and continuous membranous features visible in the overlap regions between adjacent tiles suggests both successful stitching and minimal radiation damage despite the extra dose. The diameter of each circular tile is 1.08 *μ*m.

### F. Tomogram reconstruction

Tomograms were reconstructed from the montage tiltseries using the EMAN2 (version 2.3) software package, with the projection model generated by patch-tracking [27, 28]. The tilt-series data were binned during reconstruction to a pixel size of 7.95 Å to improve contrast. Tomograms were visualized and movies generated using the 3dmod module from IMOD (version 4.10.3) [29].

## IV. Example montage cryotomograms

For proof-of-principle, we collected montage tilt-series from the thin edge of HeLa cells and reconstructed them into tomograms. During data processing, Thon rings to better than 15 Å were frequently observed on individual tiles (Fig. 2B), and distinct cellular features that spanned the overlap regions were visible throughout the tilt-series (Figs. 3 and S6). Both observations suggested that radiation damage was not severe despite the additional dose required for stitching. Consistent with this, there was no evidence of bubbling, a hallmark of radiation damage, or unevenly distributed dose in the reconstructed volumes, which were visually comparable to cryotomograms collected by standard data collection protocols. Inspection of the montage tomograms revealed rich and diverse cellular structures, including microtubules, multilamellar vesicles, mitochondria with calcium granules, actin bundles, and ribosomes (Figs. 4–6 and Movies S1-6).

**FIG. 4.**
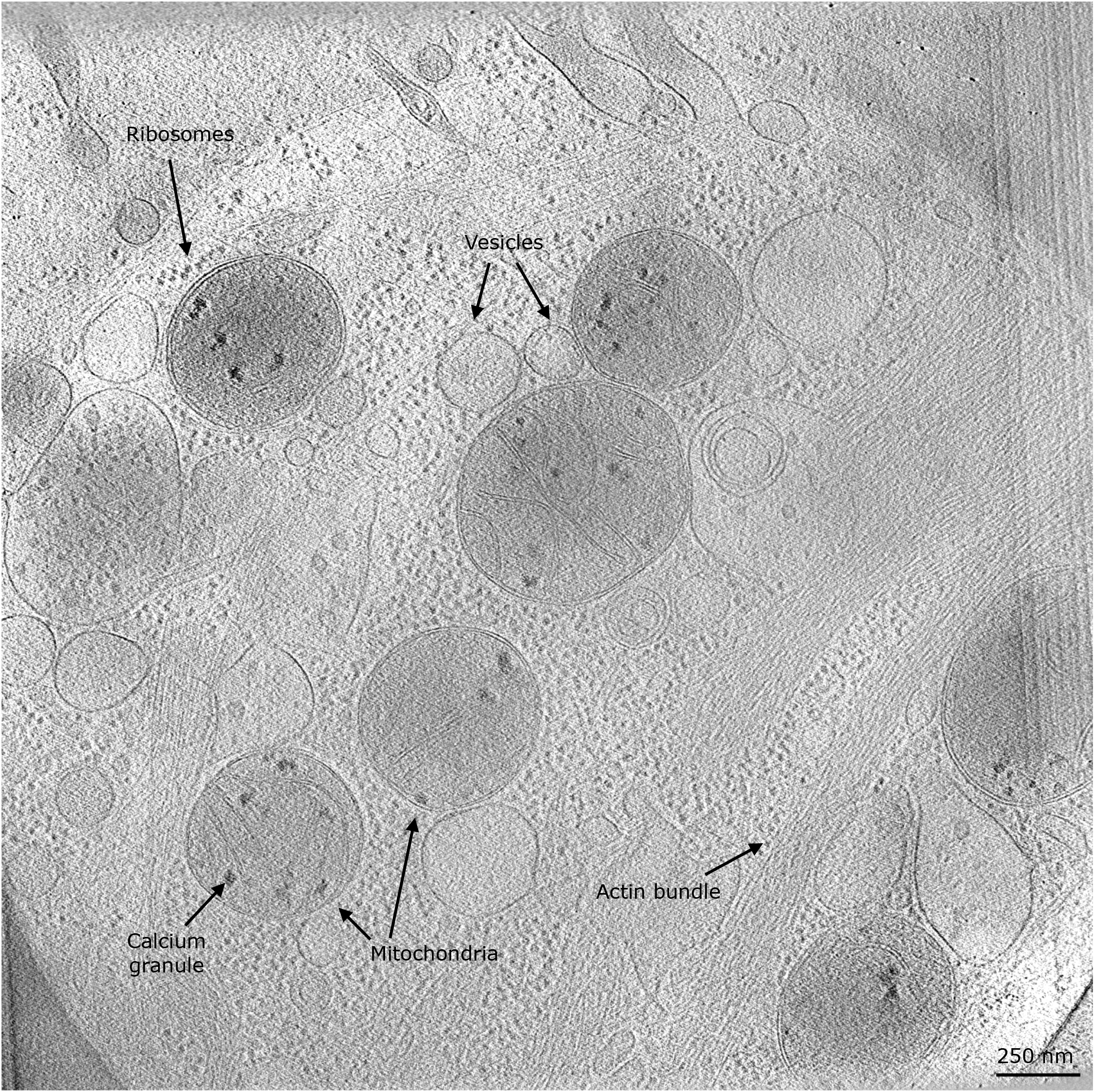
Diverse cellular features are observed in an example cryo-tomogram reconstructed from montage tilt-series. A slice is shown from a representative montage tomogram that spans a 3.3 *μ*m^2^ field of view.

**FIG. 5.**
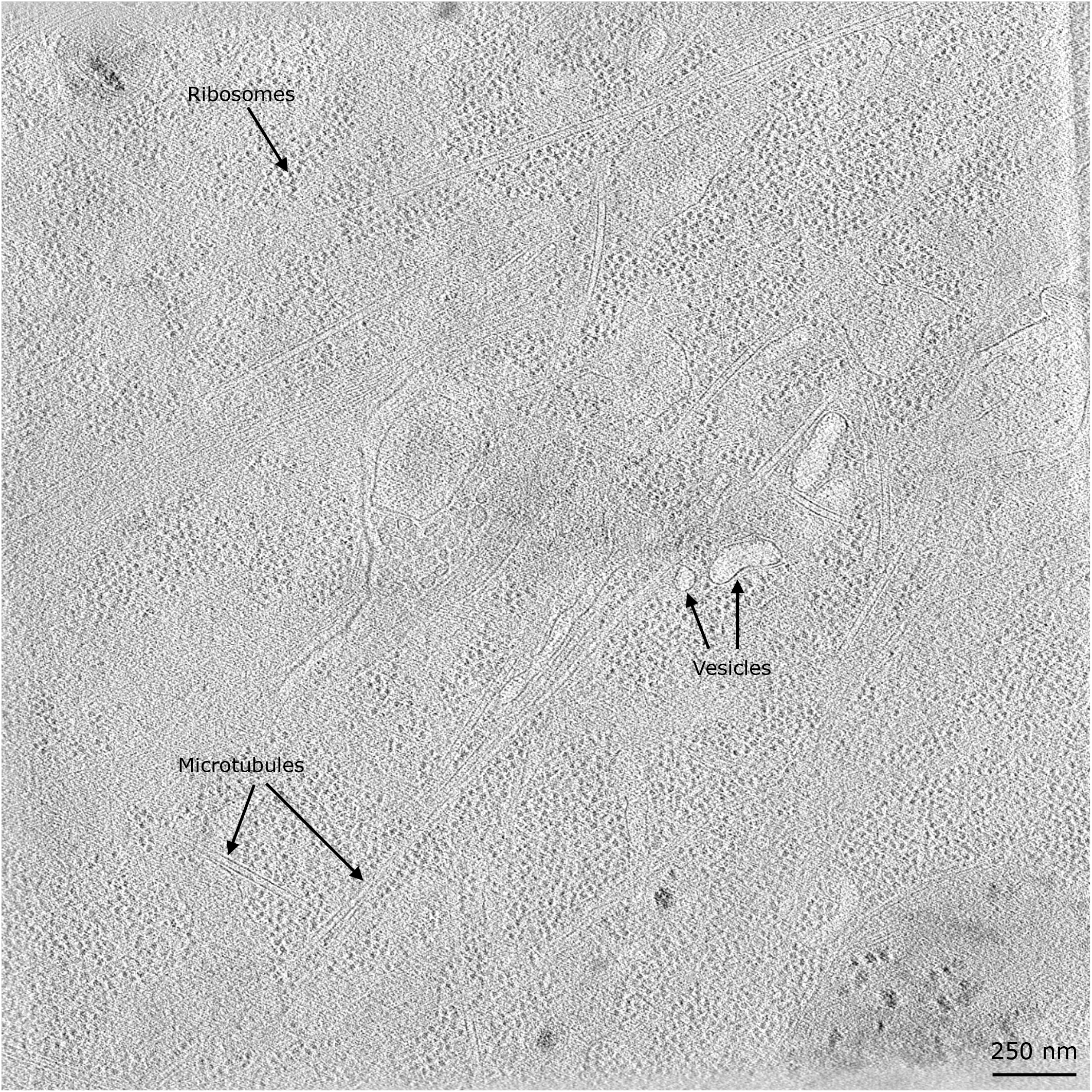
Representative montage cryo-tomogram from the thin edge of a HeLa cell. A tomographic slice spanning a 3.3 *μ*m^2^ field of view is visualized, with cellular features of interest annotated.

**FIG. 6.**
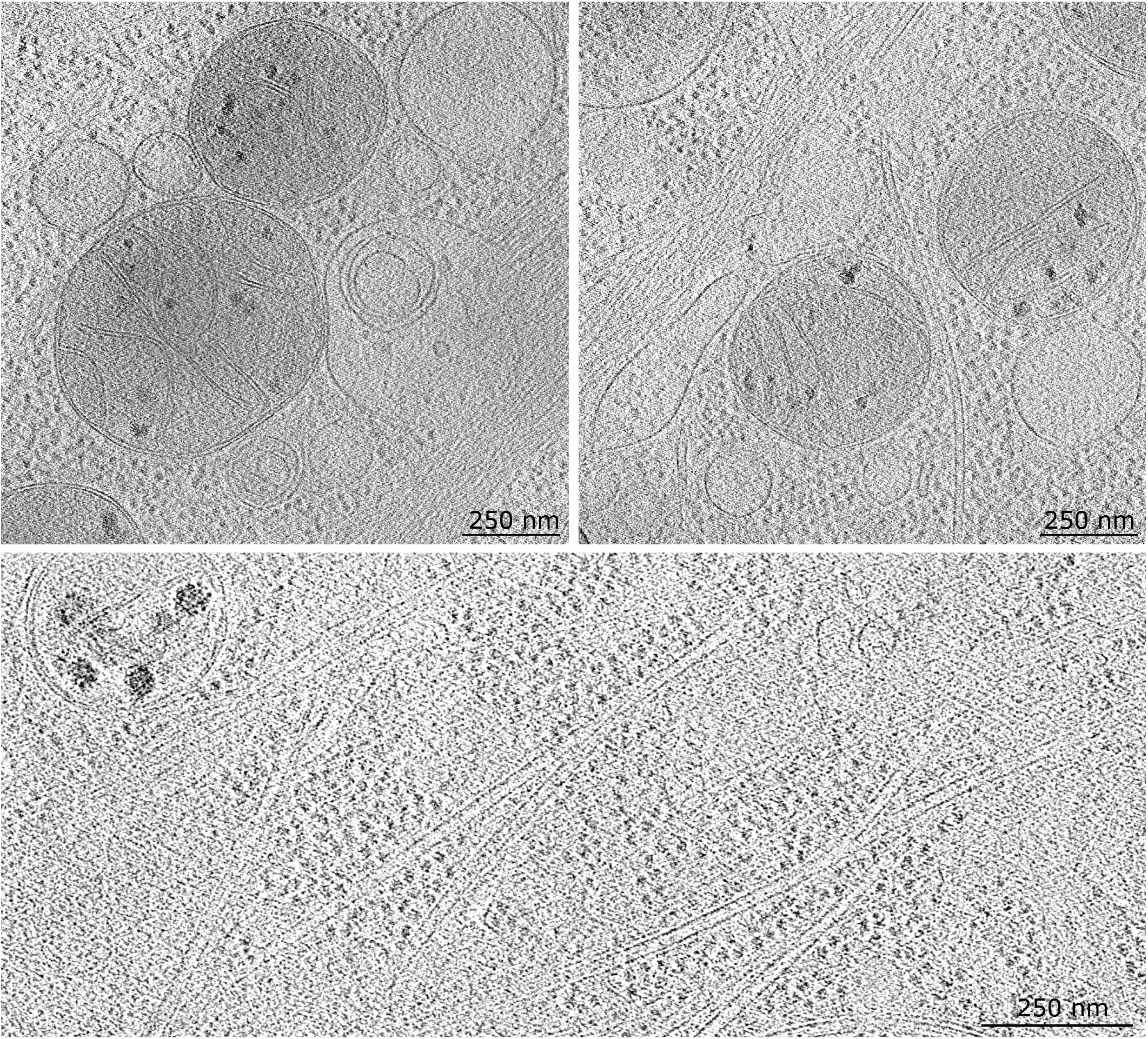
Insets from representative montage cryo-tomograms. Close-up views from the cryo-tomograms presented in Figs. 4 and 5 visualize the mitochondria (upper) and microtubules (lower) in greater detail.

## V. Discussion

Here we present a tomographic data collection and processing workflow to acquire montage tilt-series with a small pixel size but a large field of view. We used simulations to determine an acquisition strategy that efficiently distributed additional dose throughout the specimen and developed algorithms to stitch the recorded data into seamless projection images. We then assessed the efficacy of this pipeline by applying it to the thin edge of HeLa cells, yielding tomograms that spanned a 3.3 *μ*m^2^ field of view with a pixel size of 7.95 Å. The pixel size of the montage tilt-series, which would determine the Nyquist frequency of a subtomogram average, was 2.65 Å. The observation of Thon rings to better than 15 Å in the stitched images indicated that the montage tilt-series retained signal to a resolution typical of cryo-ET, despite the extra dose required for stitching that historically has been considered prohibitive to this method’s success.

Looking ahead, montage tomography will benefit from further increasing the field of view and extending the resolution of this technique. For the latter, we foresee additional ways that our data collection and processing schemes could be improved. In particular, data acquisition could benefit from collecting dose-fractionated image stacks rather than a single frame for each tile. Such “movie-mode” data collection is advantageous because it enables motion correction to reduce resolution loss due to specimen drift [30]. On the data processing side, a 3d CTF correction that spatially constrains the estimated defocus across neighboring regions may prove beneficial given the sample thickness and large variation in *z*-height of the outer tiles at high tilt angles [31, 32]. Further, additional refinement steps could be incorporated into the stitching algorithm to account for sampling warping. Correcting for this phenomenon has benefited tomogram reconstruction [3], but the magnitude and direction of this effect would be difficult to infer from projection images. An iterative approach may be possible, in which an initial 3d model of sample warping determined during tomogram reconstruction is used to refine tile registration, in turn improving the warping model and tomogram quality during a subsequent round of reconstruction.

We anticipate that montage tomography could prove particularly useful for cellular lamellae. The FIB-milling required to produce these samples is challenging and time-consuming, so maximizing the yield during data collection is a critical concern [16, 17]. By imaging a large field of view, this technique also increases the chances of capturing transient or infrequent cellular events. Finally, data collected by this method will be a valuable resource for visual proteomics, which seeks to build atlases of cellular structure at molecular resolution [33, 34]. As technical advances continue to improve the high-resolution limit of cryo-ET, montage tomography offers a way to provide more cellular context while retaining detailed views of macromolecules *in situ*.

## Data and code availability

Raw data are available at the Caltech Data Repository (https://data.caltech.edu) under accession IDs 2096, 2099, and 2103. The processed tilt-series binned to 5.3 Å can be found in the Caltech Electron Tomography Database (https://etdb.caltech.edu/). The code developed for data collection and processing is available at https://github.com/apeck12/montage.

## Acknowledgments

Data collection and analysis were respectively performed at the Beckman Institute Resource Center for Transmission Electron Microscopy and the Resnick High Performance Computing Center at Caltech. We thank Wei Zhao for preliminary samples, David Mastronarde for valuable discussions, and Tom Morrell for help with the Caltech Data Repository. A.P. is The Mark Foundation for Cancer Research Fellow of the Damon Runyon Cancer Research Foundation (DRG 2361-19). This work was supported by NIH grant AI150464 to G.J.J.

---

**FIG. S1.**
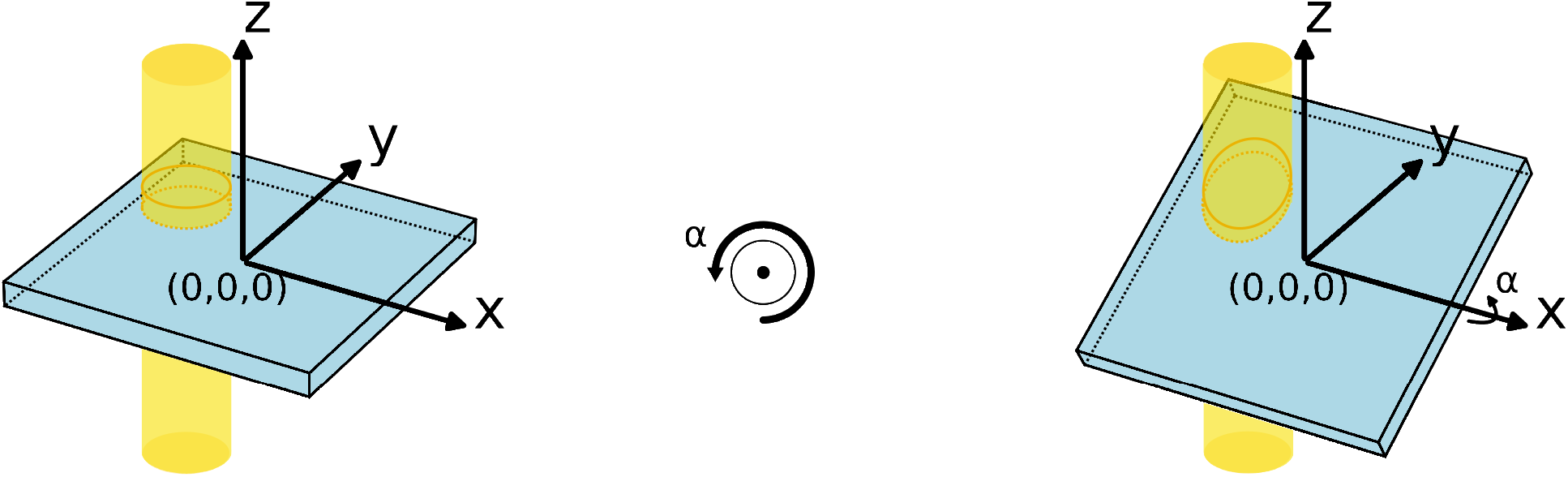
Coordinate system and axes conventions for simulating montage data collection. All simulations use the standard right-handed Cartesian coordinate system, with the origin at the center of the specimen (blue rectangular prism). The specimen is tilted around the *x*-axis. A tilt angle *α* is positive if the specimen is tilted counterclockwise when viewed from the positive side of the *x*-axis. An electron beam (yellow cylinder) is delivered to the specimen along the *z*-axis from the positive side. The illuminated volume of the specimen is the cylinder outlined by the orange ellipses. In projection the beam is circular at the first tilt angle of 0° and becomes increasingly elliptical at higher tilt angles, with the long axis of the ellipse aligned with the *y*-axis of the detector.

**FIG. S2.**
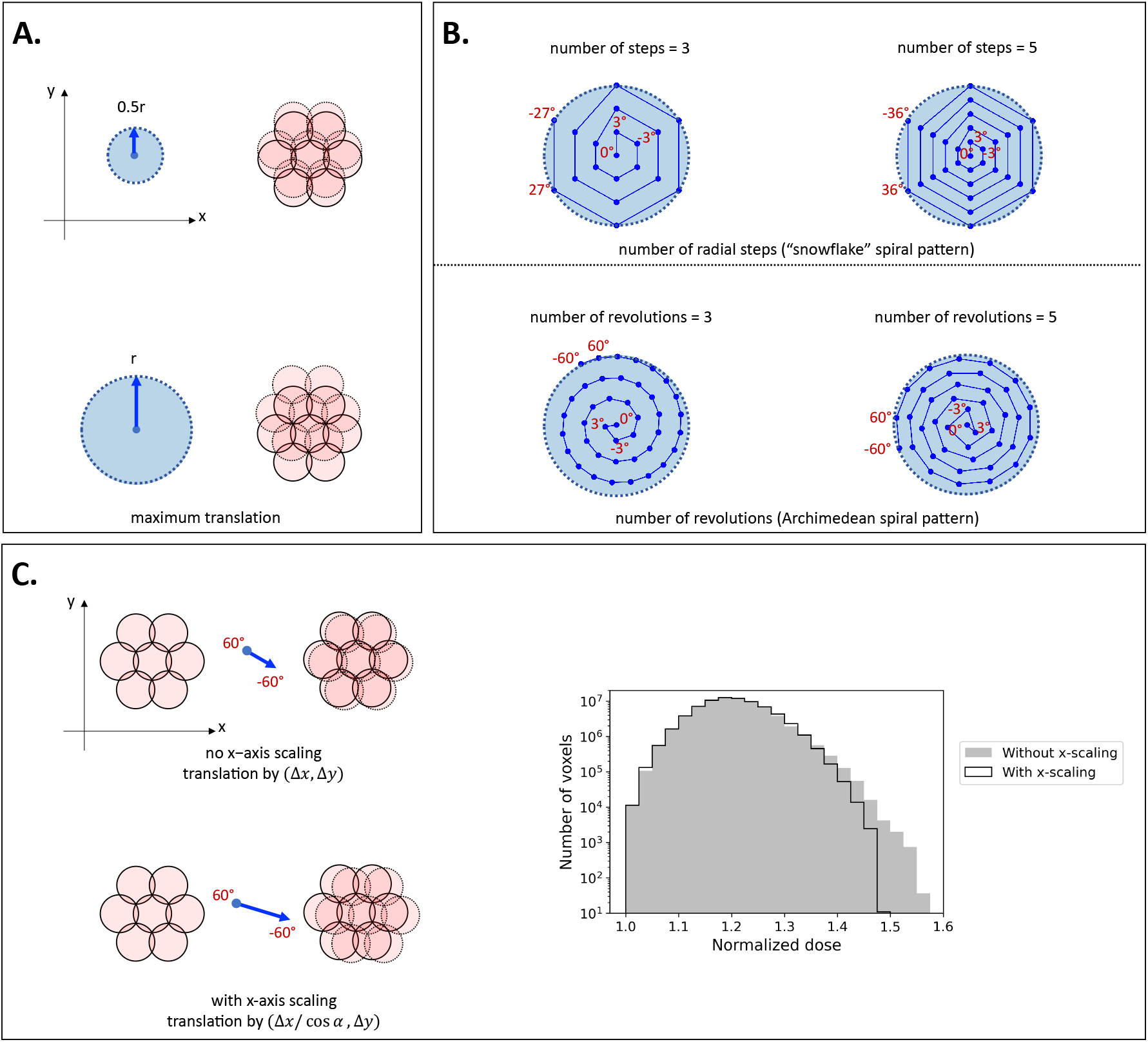
Global translational offsets of the tiling pattern change which regions of the specimen are imaged more than once at each tilt angle. For visual clarity, only the central seven tiles (pink circles) are shown. Solid and dashed borders respectively indicate tile positions before and after the indicated translation has been applied between successive tilt angles. The following parameters were varied to determine the combination of translations that optimized the efficiency of the distributed dose. (A) Maximum translation: the maximum displacement allowed for any tile during the tilt-series was varied from 0 (no offsets) to *r* (one beam radius). The blue circles on the left demarcate the region of allowed shifts for the central tile for a maximum translation of 0.5*r* (top) and *r* (bottom). The change in the overlap region for a translation of this magnitude along the *y*-axis of the detector is shown on the right. (B) Number of radial steps / number of revolutions: both parameters specify how tightly packed the spiral pattern is. Each blue dot represents the position of the central tile after the translational offset is applied at each tilt angle. The large light blue circles represent the region of allowed displacement as described in (A). For the snowflake pattern (top), the number of radial steps is the number of points between the center and edge of the light blue circle along any of the six translational directions. If the maximum translation is reached before the final tilt angle of −60°, the translational offsets for the remaining tilt angles repeat the snowflake pattern starting from the center. Archimedean spiral patterns avoid this issue by fitting all tilt angles into the light blue circles regardless of the number of revolutions (bottom). (C) Optional *x*-axis translational scaling: the *x*-axis component of all translations is optionally scaled by the cosine of the current tilt angle. The difference is subtle and most evident between tilt angles of 60° and −60°, as shown on the left, where with scaling (lower) the x-axis translational offset is twice that without scaling (upper). This scaling is motivated by the elliptical projection of the beam at high tilt angles, with the long-axis of the ellipse aligned with the *y*-axis (Fig. S1). As a result of the elliptical projections, regions of the specimen that were previously in the overlap region are more likely to be in the overlap region again along the *y*-axis than *x*-axis. To compensate for this, application of an additional *x*-axis translation that mirrored the elongation of the elliptical projection was tested. As shown by the overlaid histograms on the right, *x*-axis scaling reduced the number of voxels in the specimen that received high dose for a basic spiral pattern.

**FIG. S3.**
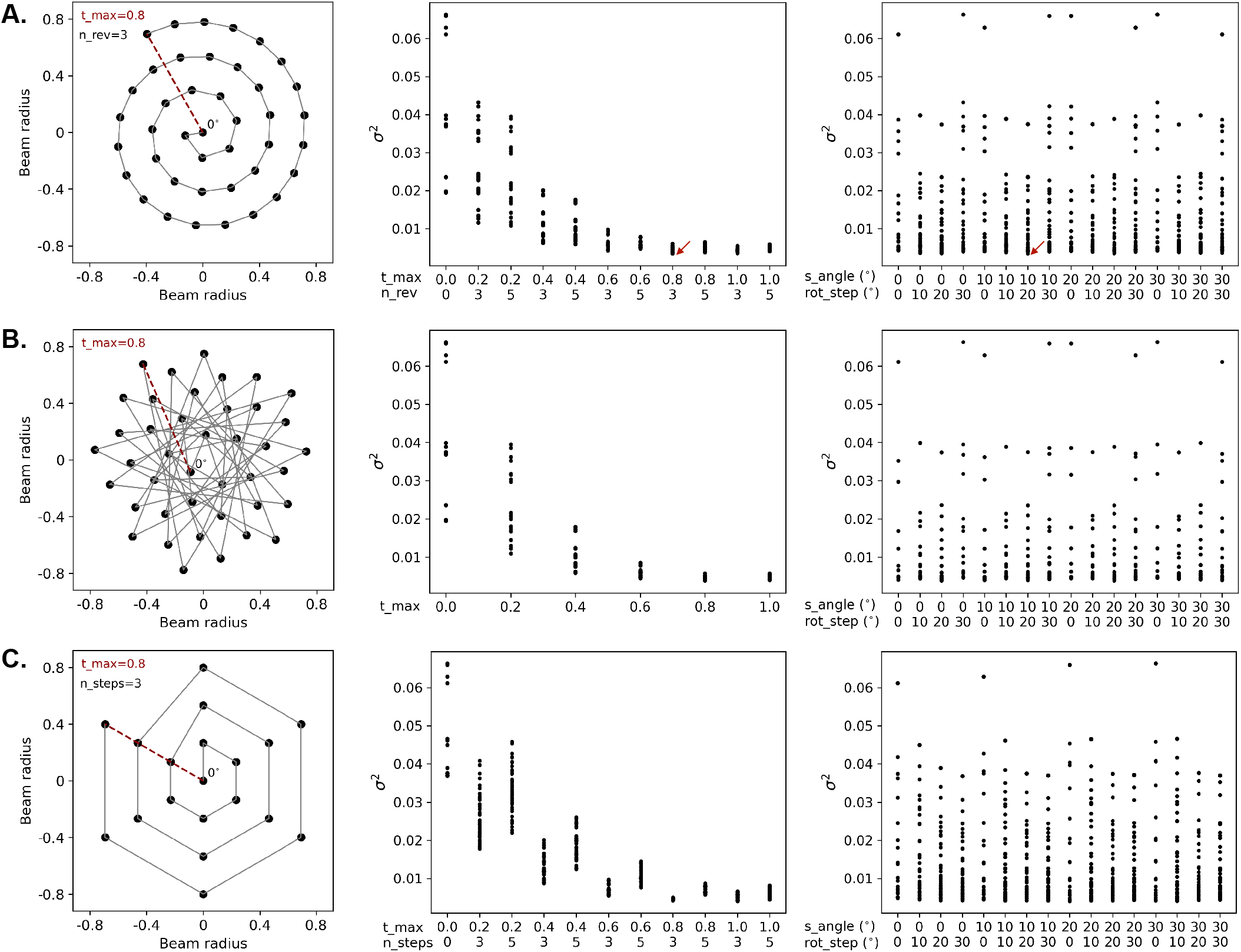
Tiling pattern efficiency is more sensitive to translational than rotational offsets between tilt angles. The changes in the position of the central tile between tilt angles for the (A) Archimedean spiral, (B) sunflower, and (C) snowflake patterns are shown at left. For each pattern, the position of the central tile at the first tilt angle of 0° is noted, and its position follows an outward spiral (indicated by the grey line) during the tilt-series. For the snowflake pattern, the original pattern is repeated after the outermost point is reached. Translational parameters are noted at the upper left. For each, Lmax (red line) indicates the maximum displacement of the central tile during the tilt-series. The parameters n_rev and n_steps correspond to the number of revolutions and radial steps for the spiral and snowflake patterns, respectively. Variants of each spiral pattern were simulated and ranked based on the variance (*σ*^2^) of the normalized dose distribution. Variance scores are shown as a function of the indicated translational parameters (center) and rotational parameters (right). For the latter, parameters s_angle and rot_step respectively refer to the initial angular offset of the hexagonally-packed tiles in the plane of the detector and the global rotation applied between tilt angles. Simulations were performed using a tilt-range of ±60° with 3° increments between tilts, applying a 1/cos exposure scheme, and assuming that Fresnel fringes contaminated 2% of the beam radius. The red arrows in (A) indicate the top-ranked pattern.

**FIG. S4.**
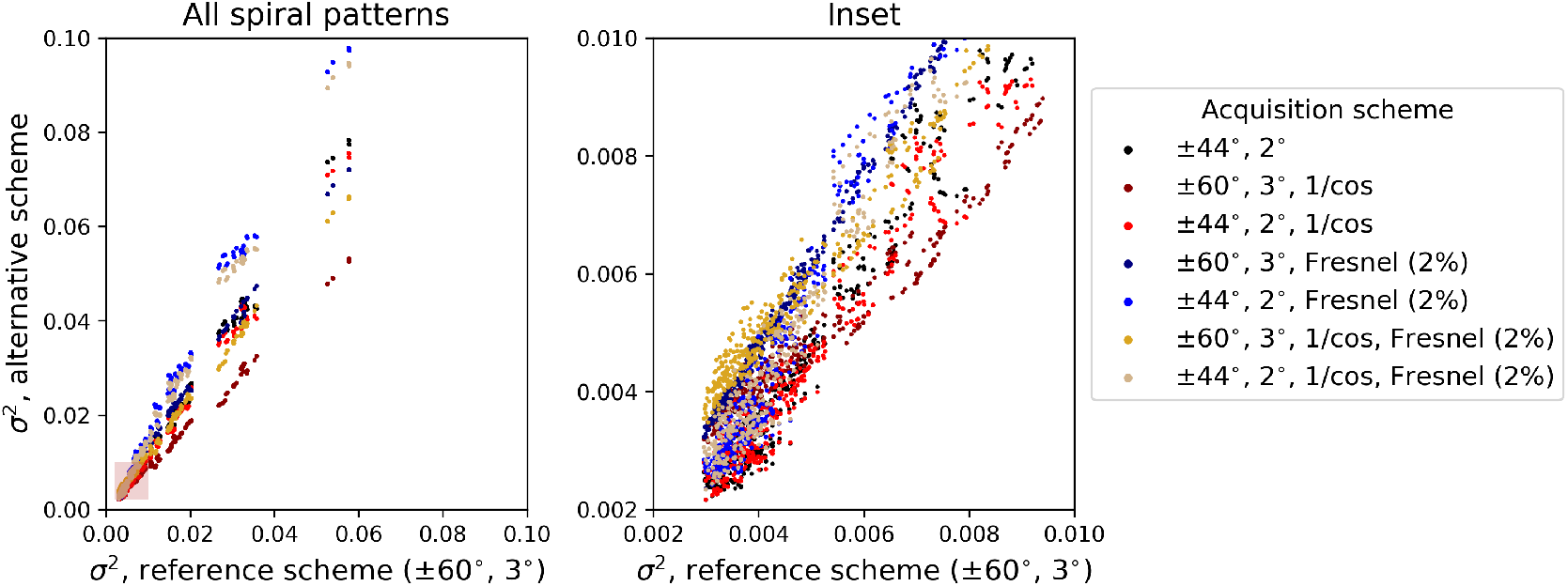
Tiling strategy efficiency is similar across different data collection schemes. 546 unique variants of the spiral pattern were tested and scored by the variance (*σ*^2^) of the accumulated dose distribution. The score of each pattern for the indicated data acquisition scheme is compared to its score for a reference data collection strategy of ±60° with 3° increments between tilt angles. Alternative acquisition schemes used a tilt-range of ±44° with 2° increments between tilt angles, increased the dose as 1/cos of the tilt-angle, and/or increased the overlap between neighboring tiles to account for Fresnel fringes spanning 2% of the beam’s radius. The transparent pink box overlaid on the highest-ranked patterns (left) is shown in the inset (right).

**FIG. S5.**
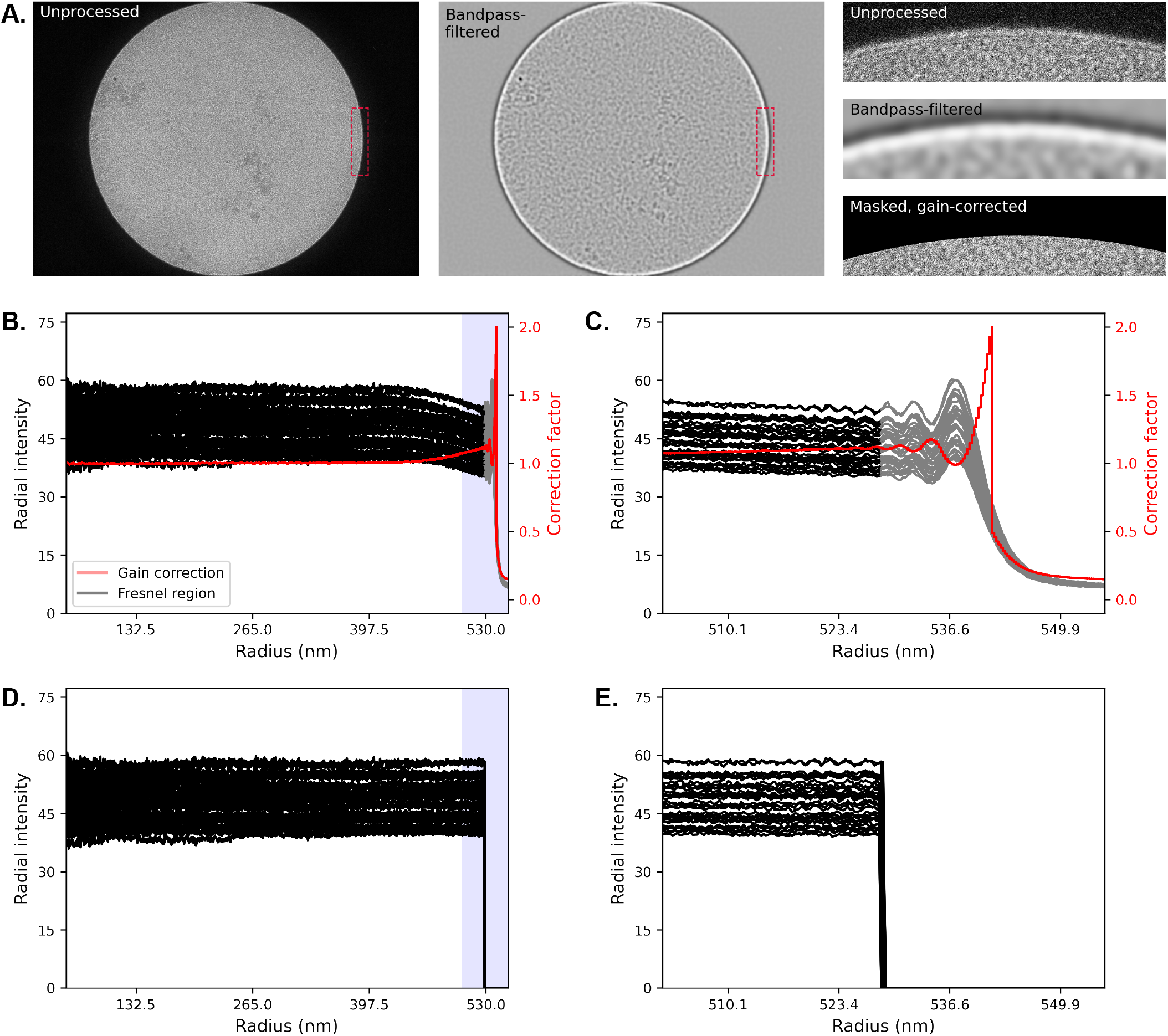
Removal of the fringe-corrupted region and correction for uneven radial illumination. (A) A representative tile is shown before (left) and after (center) applying a bandpass filter. In the filtered tile, a high intensity ring (white) coincides with the Fresnel fringes at the tile’s edge. (Right) The region boxed in red from the unprocessed, bandpass-filtered, and masked tiles visualizes elimination of the Fresnel fringes. (B) Radial intensity profiles for all 37 tiles from a representative tilt angle are plotted in black, with the region judged to be corrupted by Fresnel fringes plotted in grey. The radial gain factor used to correct uneven illumination is plotted in red. (C) Inset of the shaded blue region in (B) that focuses on the edge of the tiles affected by Fresnel fringes. (D) Radial intensity profiles of these tiles after correcting for uneven radial illumination and masking the fringes and (E) the corresponding inset.

**FIG. S6.**
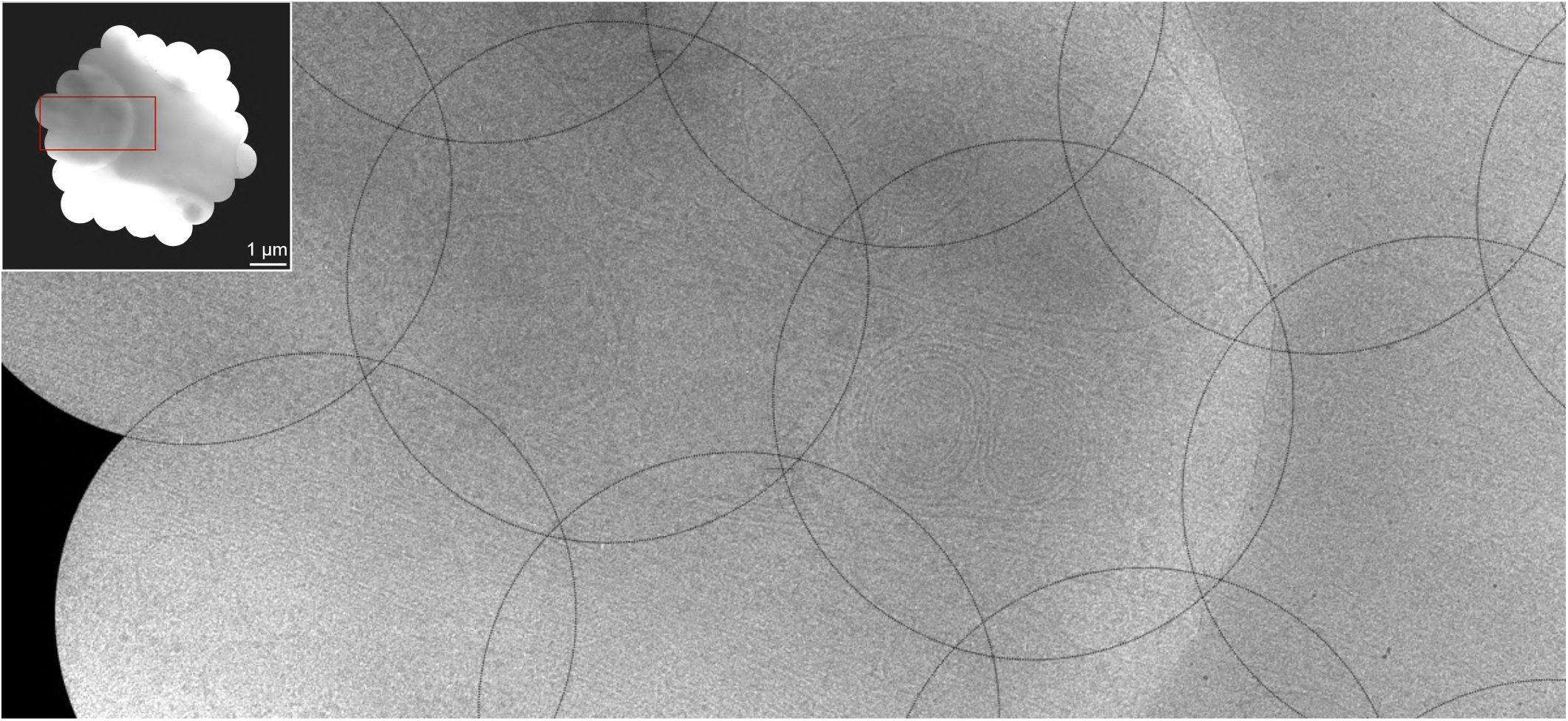
Continuous cellular features are observed in the overlap regions at intermediate tilt angles. As in Fig. 3, except the stitched tiles imaged at 30° are visualized. The inset displays the full mosaic prior to cropping for reconstruction. The region boxed in red is enlarged in the main image. The boundaries of the 1.08 *μ*m diameter circular tiles are outlined in black. The continuity of membrane features in the overlap regions between adjacent tiles suggests minimal radiation damage, even at an intermediate stage of data collection.

